# Same father, same face: deep-learning reveals paternally-derived signalling of kinship in a wild primate

**DOI:** 10.1101/810820

**Authors:** Marie JE Charpentier, Mélanie Harté, Clémence Poirotte, Jade Meric de Bellefon, Benjamin Laubi, Peter M Kappeler, Julien P Renoult

## Abstract

Animal faces convey important information such as individual health status^1^ or identity^2,3^. Human and nonhuman primates rely on highly heritable facial traits^4,5^ to recognize their kin^6–8^. However, whether these facial traits have evolved for this specific function of kin recognition remains unknown. We present the first unambiguous evidence that inter-individual facial similarity has been selected to signal kinship using a state-of-the-art artificial intelligence approach based on deep neural networks and long-term data on a natural population of nonhuman primates. The typical matrilineal society of mandrills, is characterized by an extreme male’s reproductive skew with one male generally siring the large majority of offspring born into the different matrilines each year^9^. Philopatric females are raised and live throughout their lives with familiar maternal half-sisters (MHS) but because of male’s reproductive monopolization, they also live with unfamiliar paternal half-sisters (PHS). Because kin selection predicts differentiated interactions with kin rather than nonkin^10^ and that PHS largely outnumber MHS in a mandrills’ social group, natural selection should favour mechanisms to recognize PHS. Here, we first show that PHS socially interact with each other as much as MHS do, both more than nonkin. Second, using artificial intelligence trained to recognize individual mandrills from a database of 16k portrait pictures, we demonstrate that facial similarity increases with genetic relatedness. However, PHS resemble more to each other than MHS do, despite both kin categories sharing similar degrees of genetic relatedness. We propose genomic imprinting as a plausible genetic mechanism to explain paternally-derived facial similarity among PHS selected to improve kin recognition. This study further highlights the potential of artificial intelligence to study evolutionary mechanisms driving variation between phenotypes.

## INTRODUCTION

In almost all living species, kinship is a major evolutionary force driving individuals’ relationships through kin selection processes^10,11^. Facial traits probably play a major role in recognizing kin. In humans, for example, facial traits are both the most morphologically variable and the most singular and recognizable features of the physical appearance^12^. Facial traits are also highly heritable, some showing more than 90% of genetic heritability^4,5^, resulting in elevated facial similarity among relatives, including across generations. As such, both human or non-human primate subjects scored genetic relatedness from faces at the intra-specific^6,7,13,14^ or inter-specific^15,16^ levels. However, whether kin-biased facial similarity merely reflects genetic ancestry or results from selection processes to improve kin recognition, remains unknown. Here, we test whether inter-individual facial similarity among paternal relatives has been kin-selected using a state-of-the-art artificial intelligence approach based on deep neural networks (DNNs) and long-term data obtained from a natural population of primates.

Many non-human primates show typical multimale-multifemale social groups strongly structured by matrilineal families of maternally related females associated with male-biased dispersal. In several of these societies, reproduction is skewed towards a few males, especially the alpha male, although male’s tenure does not generally last for more than a few years^17^. This general pattern is observed in mandrills (*Mandrillus sphinx*), an Old World primate living in the deep forests of Central Africa. In this species, reproduction is seasonal and male-male competition is fierce resulting in alpha male’s annual monopolization of about 70% of reproductive events in captivity^9^ and probably also in the wild. As a consequence of this high reproductive skew, most new-born infants from a given cohort are related through the paternal line but are generally born into different matrilines. While maternal half-siblings (MHS) are familiar with each other because they are raised in the same familial environment, paternal half-siblings (PHS) should not be familiar with each other because they live in different families. Yet, there is some evidence in captive juvenile mandrills that PHS have differentiated social relationships^18^ suggesting that they recognize each other as kin. In this study, we question how this peculiar social structure largely widespread across primate societies may have impacted the genotype-phenotype relationship by studying whether facial traits act as signals of kin ancestry.

We investigate the only natural population of habituated mandrills (ca. 220 individuals) for which detailed individually-based data on life-history, behaviour and demography are available since 2012. Based on patterns of reproduction in this population (Supplementary Results), we estimate that females have on average 2.1 times more PHS than MHS in their social group during the course of their life. Across these females, however, the variance in the number of PHS is 12.8 times greater than that of MHS. Selection pressures for communicative traits (facial traits, a face recognition system, or both) should thus be high because of high kin-selected benefits to recognize PHS. They are, indeed, potentially numerous in a given social group meaning that females may benefit from social support obtained from numerous kin thereby increasing inclusive fitness. Communicative traits should not be, however, error-proned because females vary a lot in their set of PHS available as potential social partners. By opposition, if reproductive skew was such that the alpha male sired all offspring, then 100% of PHS would be age-mates. With this configuration, communicative phenotypic traits allowing PHS recognition should not be selected. To sum-up, in a typical matrilineal society with potentially many PHS, individuals should recognize MHS using cues gathered from their matrilineal environment but they should also recognize PHS by relying on other cues. Given the social setting of the mandrill’s society, we hypothesize that facial traits are under strong selection to facilitate kin recognition among PHS. We first predict that PHS show strongly differentiated social relationships compared to non-kin (NK), all else being equal: they should affiliate more and be more associated than expected. Second, we predict facial traits to be under kin selection: PHS should resemble more to each other than expected given their genetic similarity. Thus, they should resemble more than NK do, but also more than MHS do, even if PHS and MHS share on average the same degree of genetic relatedness (r=0.25).

For the past 8 years, we have collated a photo-bank of c.a. 15k face pictures representing 276 individuals, some of which with portraits from birth to adulthood. This framework allowed to control for ontogenic effects on mandrill’s faces. This is particularly convenient because MHS are necessarily at least one year apart while PHS are generally age-mates. In this study, we were thus able to compare both MHS from which pictures were taken at the same age and PHS with pictures obtained at very different ages, controlling for age effects on mandrill’s faces. This unique setting would have been impossible with data collected on the short-term.

We measured similarity between female faces using DNNs. DNNs have revolutionized the computational study of facial similarity over the last five years, now outperforming human capabilities in recognizing people by their face from a photograph^20^. DNNs use a cascade of multiple layers that build representations of faces with different levels of abstraction and complexity. The deeper layers ultimately represent entire faces in an informative and low-dimensional space –the deep feature space (DFS). The DFS is informative because it is insensitive to variations that are irrelevant for the task the DNN has been trained on. For example, for face recognition a DNN learns to identify people independently of lighting, head orientation, haircut and accessories. In a DFS, two portrait images that differ only by these irrelevant variations are located at the same place, and thus the distance between images in that space reliably estimates similarity between individuals^21^. Here, we have embedded portrait images into a DFS shaped specifically to represent the identity of female mandrills. To our knowledge, this is the first study that uses a DFS to quantify phenotypic similarities in wild animals.

### Kinship and sociality

We first demonstrate behavioural biases among PHS that share more grooming behaviour and more aggression than NK, all else being equal (Table 1; Figure 1). Increased aggression probably results from increased spatial proximity among kin compared to NK that we further report (Figure 1). Surprisingly, PHS and MHS appear very similar regarding their social phenotypes which is unexpected for a matrilineal society where maternal relatedness deeply impacts the structure of female’s social relationships^22^. In captive mandrills, juveniles also show elevated affiliation both towards their adult PHS and their adult MHS; however, they affiliate more with their juvenile MHS than with their juvenile PHS probably because of highly differentiated affiliative behaviour towards a common mother at these young ages^18^. Given that adult PHS discriminate each other as kin as much as MHS do, we hypothesize that facial similarity among PHS have evolved to facilitate kin recognition.

**Table 1.** Kinship and social behaviour. Statistics obtained from Generalized Linear Mixed Models (proc GENMOD, SAS Studio) with a negative binomial distribution performed to study the relationships between social behaviour (grooming, aggression) or spatial association recorded across 45 adult females and a set of explanatory variables, including kinship.

|  | *Explanatory variables* | *Estimate* | *F* | *P* |
| --- | --- | --- | --- | --- |
| *Grooming* | Kinship<sup>1</sup> |  | 13.19 | 0.001 |
|  | MHS | 1.34 |  |  |
|  | NK | -2.67 |  |  |
|  | Rank difference<sup>2</sup> |  | 2.66 | 0.26 |
|  | 0 | -1.38 |  |  |
|  | 1 | -1.74 |  |  |
|  | Age difference | -0.02 | 0.01 | 0.92 |
| *Aggression* | Kinship<sup>1</sup> |  | 5.48 | 0.065 |
|  | MHS | -0.19 |  |  |
|  | NK | -0.98 |  |  |
|  | Rank difference<sup>2</sup> |  | 2.88 | 0.24 |
|  | 0 | 0.71 |  |  |
|  | 1 | 0.39 |  |  |
|  | Age difference | 0.19 | 3.86 | 0.050 |
| *Spatial association* | Kinship<sup>1</sup> |  | 3.35 | 0.19 |
|  | MHS | 0.39 |  |  |
|  | NK | 0.03 |  |  |
|  | Rank difference<sup>2</sup> |  | 23.23 | <0.0001 |
|  | 0 | 0.51 |  |  |
|  | 1 | 0.30 |  |  |
|  | Age difference | 0.06 | 0.14 | 0.71 |
|  | Kinship*Age difference<sup>3</sup> |  | 7.06 | 0.029 |
|  | Age*MHS | -0.09 |  |  |
|  | Age*NK | -0.13 |  |  |

<sup>1</sup>Class reference: PHS
<sup>2</sup>Class reference: 2 (2 classes of rank of difference)
<sup>3</sup>Class reference: Age\*PHS

**Fig 1.**
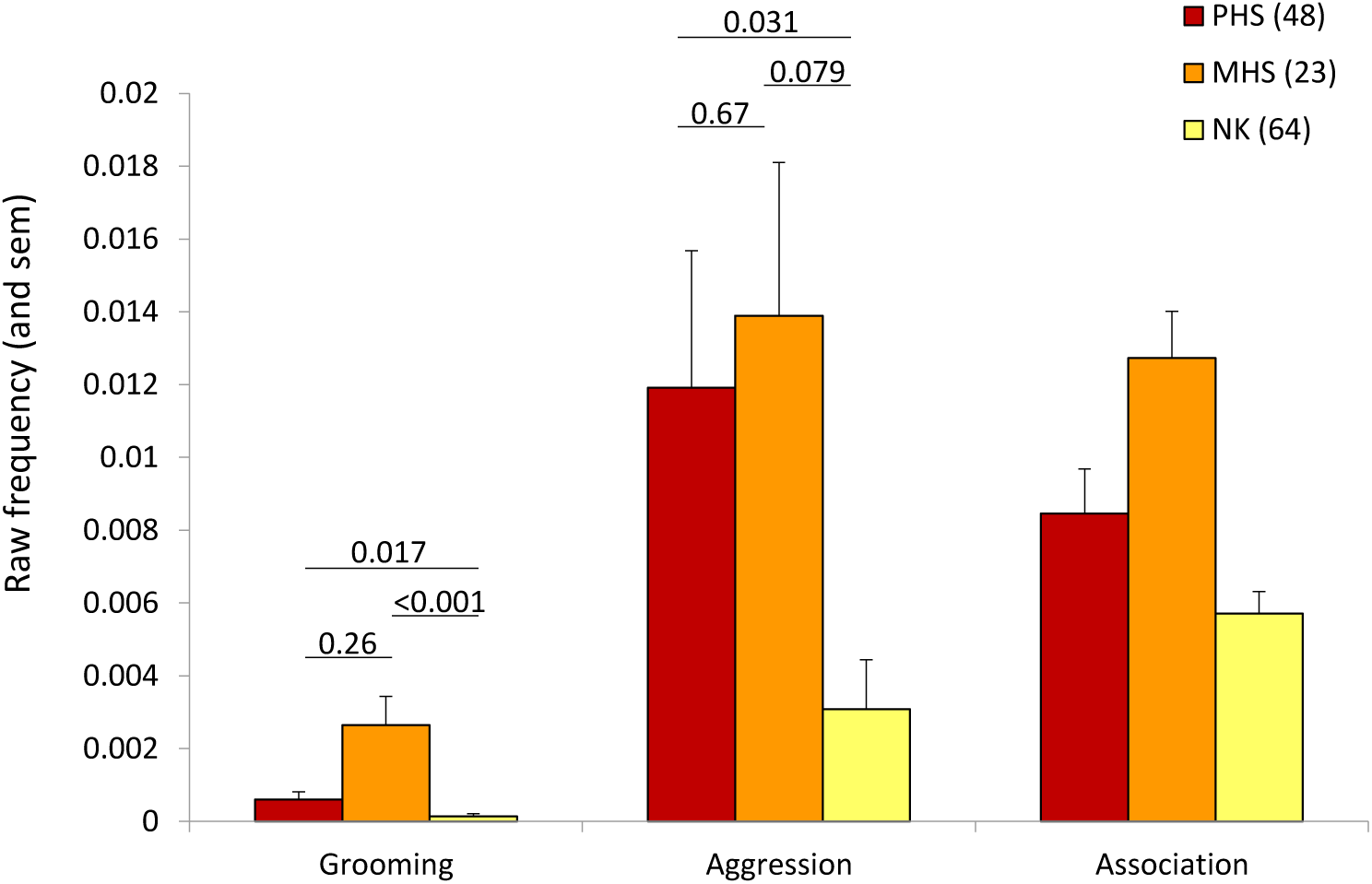
Kinship and social behaviour. Mean frequencies (and sem) of social behaviour and spatial association across kin categories composed by 45 adult females over the entire study period (2012-2019). The figure is based on raw data: time spent grooming per hour, number of aggressive behaviour observed per hour and frequency of spatial association. Pairwise differences in Least Square Means (lsmeans statement; SAS Studio) were calculated across kin categories for grooming and aggression. For spatial association, the overall effect of kinship was combined to females’ age difference. However, a closer examination of the data showed that the same general pattern as the one depicted on the figure was observed across different categories of age difference. Sample sizes are provided within parentheses.

### Genetic and facial similarities

We first studied the relationship between genetic relatedness, obtained from a well-resolved pedigree, and facial similarity across 38,515 pairs of pictures collected on 38 adult females (703 different pairs in total). While controlling for the identity of females’ pairs (random effect) and the difference in age across these pairs, we demonstrate a negative relationship between genetic relatedness and the distance between faces within the DFS (‘face distance’ hereafter; General Linear Mixed Models –LMM; estimate=-2.33, F=43.09, P<0.0001): related females look more alike than unrelated females (Figure 2). For example, compared to the average distance between faces of the same individual, the average face distance increases by 26% for pairs of individuals with a coefficient of relatedness r=0.5 corresponding to e.g. mother-offspring pairs, and by 36% for pairs sharing only r=0.0625, corresponding to e.g. simple first cousins.

**Fig 2.**
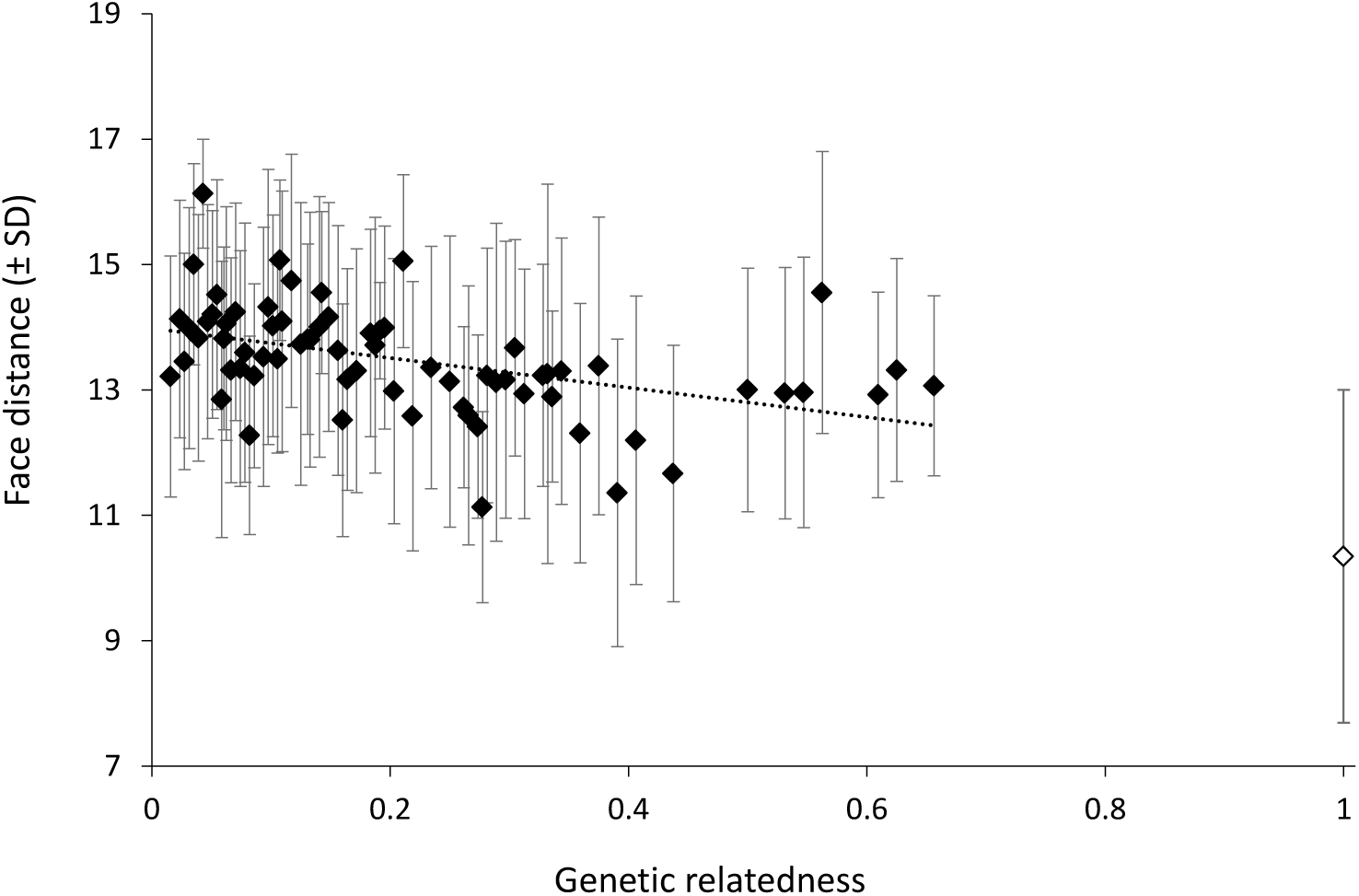
Genetic relatedness and face distance. Mean raw face distances (± SD) across the different values of relatedness observed 38 across adult females. For scaling, we have represented the averaged face distance obtained from pictures taken on the same females (i.e. intra-individual similarity). For the sake of clarity and for illustrative purposes, we have represented a regression line based on these averaged relatedness values. SD are represented rather than sem to depict the full range of variation of face distances.

### Kinship, facial similarity and female’s development

We then explored whether kinship, treated as a discrete variable, was detectable from faces and if so, when this occurred during females’ development. We restricted our analyses to MHS and PHS with an equivalent degree of genetic relatedness (r~0.25) that we compared to non-kin pairs (NK). Across 6,992 pairs of pictures collected on 45 adult females (159 different pairs in total), we show that face distance significantly differs across kin categories. This effect interacts, however, with the age difference observed across pairs of females’ pictures (Table 2). A closer examination across various age categories reveals that this interaction disappears when splitting pairs of pictures of females aged less than 2 years apart from those aged more than two years apart. Across the former pairs, PHS show the lowest face distance, i.e. they are more similar to each other than either NK or MHS are (Table 2a; Figure 3). Across the later pairs, kinship only marginally explains face distance (Table 2a), with NK being the most dissimilar compared to the two other kin categories (Figure 3). In juvenile females, we observed similar effects of kinship on face distance, in combination with age difference. As before, a closer examination of the data revealed that age distance across pairs does not impact face distance anymore when splitting pictures of juveniles aged less than a year apart from those aged more than a year apart (Table 2b). In these females very close in age and as in adults, PHS are more similar to each other than MHS or NK are. Surprisingly, MHS and NK do not differ significantly. This pattern holds true when considering pairs with more than a year apart, although kinship overall does not significantly impact face distance anymore (Table 2b; Figure 3).

**Table 2.** Kinship and face distance. Statistics obtained from General Linear Mixed Models (proc GLIMMIX, SAS Studio) performed to study the relationships between face distance and a set of explanatory variables, including kinship, in **a.** all adult female-female pairs of pictures, pairs aged less than 2 years apart and pairs aged more than 2 years apart; and in **b.** all juvenile female-female pairs, pairs aged less than a year apart and pairs aged more than a year apart.

**a.**
|  | *Explanatory variables* | *Estimate* | *F* | *P* |
| --- | --- | --- | --- | --- |
| *All pairs*<br>N = 6992 pairs, 45 females | Kinship<sup>1</sup> |  | 18.18 | <0.0001 |
|  | MHS | 0.51 |  |  |
|  | NK | 1.21 |  |  |
|  | Age difference | 0.04 | 3.07 | 0.080 |
|  | Kinship*Age difference<sup>2</sup> |  | 9.13 | 0.0001 |
|  | Age*MHS | -0.08 |  |  |
|  | Age*NK | -0.11 |  |  |
| *In females ≤ 2 yrs*<br>N = 2421 pairs, 43 females | Kinship<sup>1</sup> |  | 16.48 | <0.0001 |
|  | MHS | 0.79 |  |  |
|  | NK | 1.43 |  |  |
|  | Age difference | -0.17 | 4.86 | 0.017 |
| *In females > 2 yrs*<br>N = 4571 pairs, 44 females | Kinship<sup>1</sup> |  | 2.96 | 0.052 |
|  | MHS | -0.37 |  |  |
|  | NK | 0.27 |  |  |
|  | Age difference | -0.02 | 0.97 | 0.32 |

<sup>1</sup>Class reference: PHS
<sup>2</sup>Class reference: Age\*PHS

**b.**
|  | *Explanatory variables* | *Estimate* | *F* | *P* |
| --- | --- | --- | --- | --- |
| *All pairs*<br>N = 1589 pairs, 16 females | Kinship<sup>1</sup> |  | 12.04 | <0.0001 |
|  | MHS | -2.97 |  |  |
|  | NK | 2.66 |  |  |
|  | Age difference | -0.28 | 18.40 | <0.0001 |
|  | Kinship*Age difference<sup>2</sup> |  | 31.93 | <0.0001 |
|  | Age*MHS | 7.78 |  |  |
|  | Age*NK | -1.49 |  |  |
| *In juveniles ≤ 1 yr*<br>N = 805 pairs, 16 females | Kinship<sup>1</sup> |  | 3.85 | 0.022 |
|  | MHS | 2.42 |  |  |
|  | NK | 1.66 |  |  |
|  | Age difference | -0.59 | 2.08 | 0.15 |
| *In juveniles > 1 yr*<br>N = 784 pairs, 13 females | Kinship<sup>1</sup> |  | 0.92 | 0.40 |
|  | MHS | 2.97 |  |  |
|  | NK | 0.88 |  |  |
|  | Age difference | -2.62 | 39.97 | <0.0001 |

<sup>1</sup>Class reference: PHS
<sup>2</sup>Class reference: Age\*PHS

**Fig 3.**
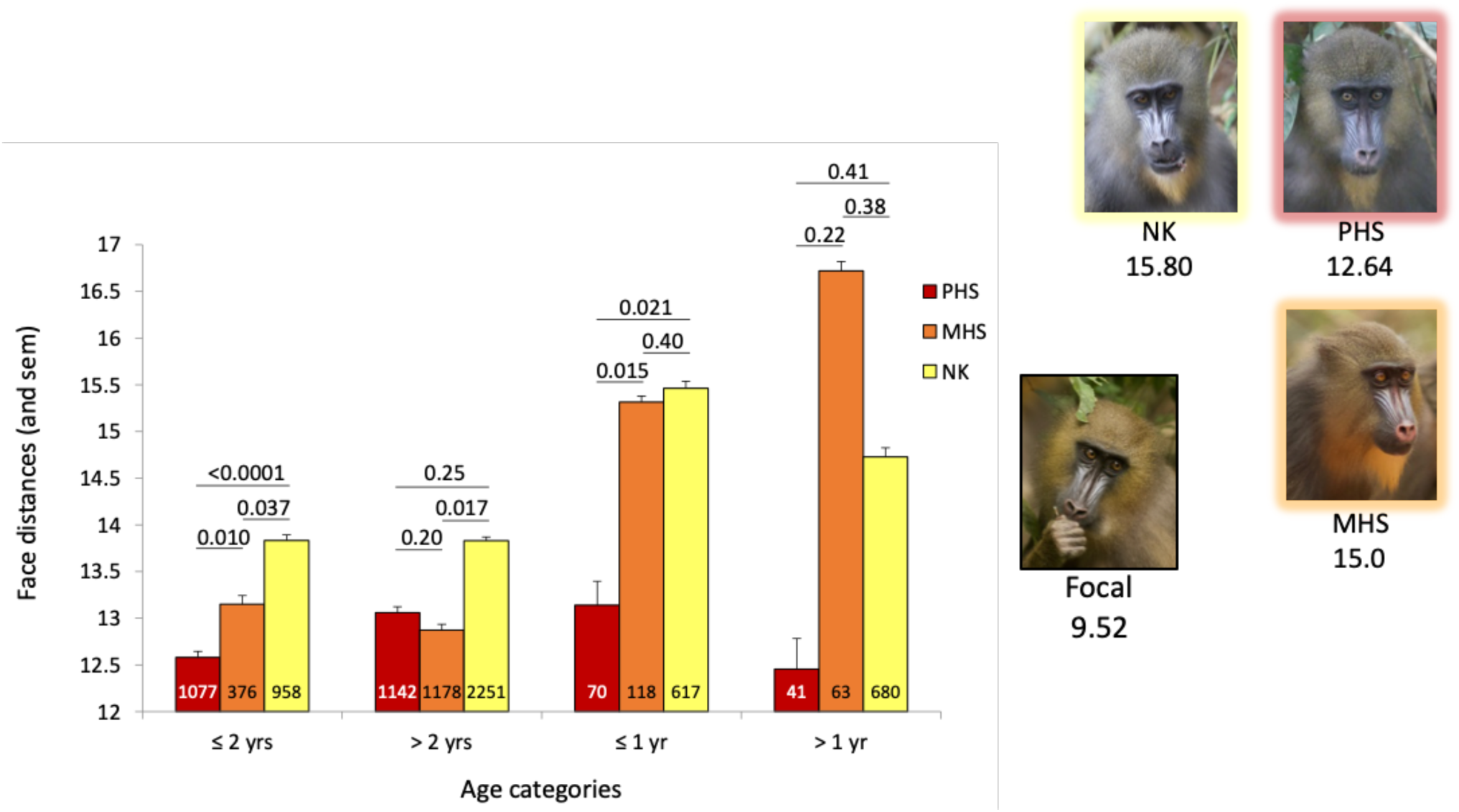
Kinship and face distance. Mean raw face distances (and sem) across kin categories for pictures taken on 45 adult and 16 juvenile females at two age differences. Pairwise differences in Least Square Means (lsmeans statement; SAS Studio) were calculated across kin.age categories. Sample sizes are provided within bars.

If individuals tend to interact more with paternal than maternal relatives, the kinship theory of genomic imprinting predicts a differential expression of patrigenic *vs*. matrigenic alleles, such as those related to facial traits, because of different consequences for their respective inclusive fitness^23^. We propose that this molecular mechanism may explain our results. Because of their complex social organization, mandrills would strongly benefit from recognizing the numerous PHS available for socializing or forming coalitions with. Alternatively, the greater facial similarity among PHS may be a by-product of an exaggerated resemblance to a common father that may have evolved to facilitate e.g. paternal care in this promiscuous primate. In a polygynous human population, for example, paternal investment is positively correlated to father-child facial similarity^24^. We think, however, that this scenario does not hold true for our study system because i) mandrills are highly dimorphic and an adult male does not resemble to any other mandrills (Supplementary Figure 1), ii) males are only temporary resident in the social group of their offspring, and iii) evidence of true paternal care are limited in this species^18^. In humans, genomic imprinting has been proposed to account for the temporary greater mother-infant facial similarity compared to fathers through pleiotropic effects. Indeed, there is a preferential expression of maternal genes during foetal development to control resource allocation^25^. Mother-child early facial similarity would be a by-product of genomic imprinting during the foetal life, explaining why similarity to mothers decreases between birth and one year of age^26^. By contrast, in mandrills, the greater facial similarity among PHS comes early in age but also lasts until adulthood and long after their father’s secondary dispersal from their group. This difference between humans and mandrills may originate from the distinct social organization of mandrills. Females may have numerous PHS in their group throughout their life; however, they live in different maternal families. This social setting has probably generated intense selection on communicative traits allowing kin recognition. Accordingly, we recently found that genetic relatedness was also encoded into mandrills’ voice^27^. These phenotypic cues of relatedness produce differentiated behavioural responses, as captive mandrills are able to discriminate unfamiliar relatives based either on acoustic cues^27^ or on visual cues^6^ alone.

These results leave us with an open question about the ability of female mandrills to detect their own facial similarity to others. Although there are evidence that some primates know what they look alike^28^, a more plausible explanation is that association and behavioural biases among PHS are mediated by third parties, such as mothers. If mothers evaluate the facial similarity of their offspring during juvenescence and behave differentially accordingly, then behavioural biases may pertain in juveniles until adulthood. The fact that the difference in facial similarity between MHS and PHS is particularly pronounced in juvenile females and that MHS are indistinguishable from NK at these young ages support this view. Collecting detailed behavioural data on mothers’ associations will eventually reveal such a pattern, highlighting again the value of long-term field projects to generate such inestimable but long-lasting data^29^.

This study highlights the potential of deep learning for the quantitative study of complex animal phenotypes. Faces are a typical example of such phenotypes: they can be described in multiple ways, emphasizing either details like skin texture or the specific position of a mole, or more global features like the contour roundness^30^. Describing faces has been therefore an historical challenge for both computer scientists and psychologists. The once popular eigenface approach could summarize all those dimensions^31,32^ but it suffered from excessive sensitivity to noisy variation (e.g., lighting). In the 2000s, handcrafted local and noise-tolerant descriptors (e.g. Gabor filters^33^) then became the dominant approaches, but they required too many descriptors to capture the whole complexity of faces. The explosive popularity of DNNs in face recognition studies is explained by their capacity to circumvent these limitations altogether^34^. But beyond faces, DNNs can measure similarity between any kinds of complex phenotypes. Recently, Cuthill and colleagues^35^ used distances within a DFS to investigate mimicry in butterflies from museum specimens. With our study, we highlight that a DFS further allows to investigate animal phenotypes in field conditions where standardizing animal’s position is impossible.

## METHODS

### Studied population

Since April 2012, we are monitoring the only habituated social group of free-ranging mandrills worldwide, inhabiting the Lékédi Park in Southern Gabon (Bakoumba) within the framework of the “Mandrillus Project” (www.projetmandrillus.com). This group originates from 65 captive-born mandrills housed at CIRMF (Centre International de Recherches Médicales de Franceville, Gabon) and released into the park on two occasions (36 individuals released in 2002 and 29 in 2006; see for details^36^). At the end of this study (June 2019), the group was composed of ca. 220 individuals of both sexes and all ages, about 160 of them being individually-known and daily monitored^37,38^. During everyday observations, we typically record detailed data on group living and composition as well as on social behavior. We studied a total of 50 females that contributed to one, two or to the three datasets as follows: 16 juvenile females aged 1.3-3.8 and 45 adult females aged 4.1-21.3 at the time of photograph collection and 45 adult females aged 4.4-26.4 yrs at the end of the collection of social behavior (June 2019). Dates of birth were either known to a few days thanks to daily monitoring for 28 females or approximated using general condition and patterns of tooth wear for the remaining 22 females^39^. For most of these later females (70%), the error made in age estimation was less than a year. Removing from our analyses the few females for which the error made was estimated to be more than a year did not change our results (not shown).

### Genetic relatedness and kinship

DNA from the 50 studied females was extracted from either blood (46 females) or fecal (4 females) samples. Blood was collected during annual trappings that occurred from 2012 to 2015 (see for details^6^). Fecal samples were collected on each occasion. DNA extractions from either the buffy coat or from fresh fecal pellet were performed using resp. QIAamp DNA Blood or stool Mini Kits (Hilden, Germany). Microsatellite genotyping was carried out using 12-36 primer pairs^40,41^. Paternity analyses were performed using Cervus 3.0 software using previously described procedures^41^. We reconstructed the full pedigree of the 16 females born in captivity going back as far as the generation of unrelated founder animals^40,41^. We genetically determined both parents for 25 individuals out of the 34 individuals born into the wild. Pairwise genetic relatedness was calculated from the pedigree only for those individuals with at least the four parents unambiguously known, using ENDOG v4.8^42^. As well, only these pairs of females with the four parents known served as possible NK. For the remaining nine females, we knew only the mother’s identity (eight females) or the father’s identity (one female) because the genetic sample did not match any adult male or female of the genetic database. We used these nine females to determine PHS or MHS pairs because none of these resulting pairs may have been full-sibs. In addition to PHS (sharing the same father) and MHS (sharing the same mother) pairs, we considered as NK these females that shared less than r ≤ 0.0325. We excluded the few full-sibs from our data sets.

### Behavioural observations

Since August 2012, trained observers, blind to the protocols involved in this study, performed behavioural observations on 45 adult females (≥ 4 yrs) using 5-min focal sampling (1776 hours of focals total, mean per female ± SD: 39.5 ± 41.8). During these focals, all social interactions between these females, including time spent grooming and aggressive behaviour (grasp, bite, chase, lunge, ground-slap, head-bob) considered as bouts, were recorded. In addition to grooming and aggression, we considered spatial associations: three times during each focal, we scanned and recorded all studied females located less than 5 meters away from the focal female. We pooled all the behavioural data over the entire study period to improve our statistical models (see below). For these behavioural analyses, we considered 48 pairs of PHS, 23 pairs of MHS, and 64 pairs of NK for which the age difference was ≤ 6.5 yrs because, in this dataset, there was no PHS more different in age that this threshold.

Female dominance rank was evaluated using the outcomes of approach-avoidance interactions collected during focals or *ad libitum* observations and calculated using normalized David’s score (as per^43^). We divided adult females into three categories of rank of similar size across the entire study period (high-ranking, medium-ranking, low-ranking).

### Measuring facial similarity

We estimated the similarity between female mandrills through the distance of their portrait images in a DFS. We used a two-step pipeline that performs a face identification task followed by a face verification task (Figure 4). In the face identification task, a DNN is trained to identify mandrills, thereby learning a DFS adapted to represent mandrill faces. In the face verification task, a distance metric is learned such that the distance between same-individual pictures in the DFS is minimized. This metric is eventually used to predict the similarity between faces in the studied population of mandrills. The pipeline has been developed in MATLAB; methods and results are detailed in the electronic Supplementary Information.

**Figure 4.**
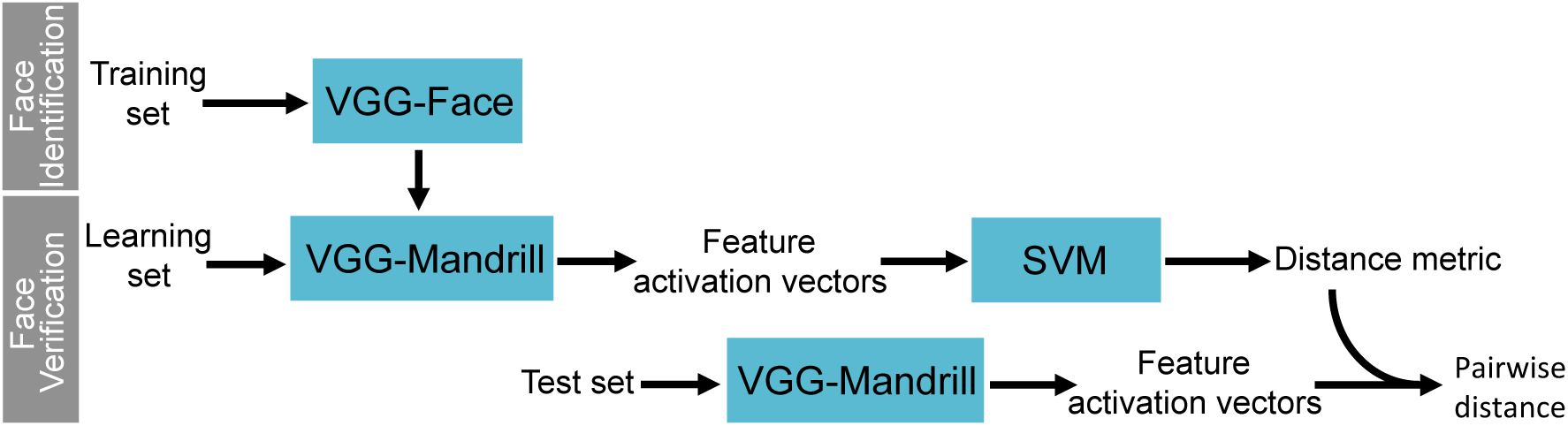
Pipeline for measuring facial similarity. The pipeline contains two main steps. During the face identification step, a deep neural network previously trained for human face recognition (VGG-Face) is retrained to identify mandrill faces. The newly trained network is then used in a face verification task, first to learn a distance metric using a Support Vector Machine (SVM) trained to detect whether two faces represented by their feature activation vectors (i.e. coordinates in the deep feature space) represent the same individual or not, then to compute the resemblance (i.e. the distance in the deep feature space) between pairs of portrait images for the studied population.

#### Image datasets

Our database includes ~16k portrait images of 276 different mandrills. This database was split into a learning set and a test set, which are different for the adult female and juvenile analyses (Supplementary Methods). For the adult female analysis, the learning set included pictures of semi-captive and captive males and females of all age classes (obtained from captive groups), as well as wild individuals from the studied population, except adult females. For the juvenile analysis, the learning set was the same as above but it included adult females from the studied population but excluded juveniles from this population. Because the face of a given individual varies considerably between its different age categories (Supplementary Results), we trained the algorithm to recognize ind.age categories (i.e. individual at a given age category) rather than individual classes. The learning sets were in turn split into a validation set (2 images per category) and a training set (other images). The validation set was used to compare the accuracy of identification between different parameter settings; however, for measuring similarity in the test set, we used a DNN trained on the entire learning set. We compared training sets with different levels of image quality and number of images per class (Supplementary Methods). The test set included either adult or juvenile females from the study population. All portraits were downsized to 224×224 prior to analyses.

#### Embedding portrait images into a deep feature space

We used the popular VGG16 architecture to recognize mandrills individually. Rather than training the algorithm from scratch (i.e., initializing it with random weights), we used VGG-Face^20^ as a starting point and retrained this network with mandrill portraits. This procedure, called “transfer learning”, allows to reach high model performance even with relatively small datasets^44^. VGG-face is a VGG16 that previously learned to recognize 2,6k different humans from a total of 2,6M portrait pictures. As typical with DNNs, VGG-Face builds representations of faces hierarchically: its first, shallow layers represent very simple and local features, for example skin colour and texture; the medium layers combine these features to represent more complex shapes and colour patterns like a pupil; the deeper layers combine previous features to represent an entire face; and the very last layer eventually classifies face images into different ind.age classes^34^. Transfer learning exploits the fact that features of shallow layers usually describe universal properties of images, and thus need no or minimal tuning when used for a new task or with a new dataset^34,44^, contrary to deep features that are task- and image-specific. For fine-tuning VGG-Face with mandrill images, we thus set a very small (10^−5^) learning rate for shallow and medium layers, a slightly larger learning rate for deep layers (10^−3^, decreasing down to 10^−5^ at the end of training) and we entirely replaced the last, classification layer (Supplementary Methods). After approximately 15 epochs, VGG-Mandrill, our newly trained DNN, was able to recognize individuals with high accuracy: 91.1% and 91.9% of correct identification for the adult female and juvenile validation set, respectively. This high performance indicates that VGG-Mandrill was able to build deep feature space (DFS) that informatively represents mandrill faces.

#### Distance metric learning

Facial similarity can be estimated by simply measuring the distance between images embedded in the DFS. However, not all features defining the DFS are similarly good for estimating facial similarity. Following the method previously developed^21^, we learned a similarity metric that calculates the weights of features that optimize a face verification task. We first extracted the feature activation vector (i.e. the coordinates in the DFS) of all images of the training set in the DFS of VGG-Mandrill. Next, we randomly selected 15k pairs of images representing different individuals and 15k pairs representing same individuals, and for each of these pairs we calculated the *χ*^2^ difference (*f*_1_[*i*]-*f*_2_[*i*])^2^/(*f*_1_[*i*]+*f*_2_[*i*]), where *f*_1_ and *f*_2_ are the feature activation vectors of the two images and *i* the feature index (the *χ*^2^ difference has the same dimensionality as feature activation vectors). Then, we run a linear support vector machine (SVM) with the *χ*^2^ difference vector in explanatory variables and 0 (different-individual pairs) or 1 (same-individual pairs) as a response variable. This SVM output feature weights *ω*_i_, which were used to calculate a weighted *χ*^2^ distance as *χ*^2^(*f*_1_,*f*_2_) = Σ_i_*ω*_i_(*f*_1_[*i*]-*f*_2_[*i*])^2^/(*f*_1_[*i*]+*f*_2_[*i*]). When evaluated with test sets, the SVM-based face verification model was able to identify whether two pictures represented the same individual with an accuracy of 83% for the adult female set and 90% for the juvenile set (Supplementary Methods). We calculated the weighted *χ*^2^ distance between all pairs of images in the test sets.

#### Studied pairs and validation

To study facial similarity in adult females, we considered 50 pairs of PHS, 30 pairs of MHS, and 79 pairs of NK for which the age difference was ≤ 9.3 yrs because, in this dataset, there was no PHS more different in age that this threshold. In juvenile females, we studied 16 pairs of PHS, 5 pairs of MHS, and 27 pairs of NK for which the age difference was ≤ 1.8 yrs for the same reason. We checked that intra-individual facial similarity (identical female with photographs taken at various ages) was greater than any other pairs analysed (Supplementary Table 1).

### Statistical analyses

#### Kinship and sociality

To study the relationship between kinship and social behaviour, we summed-up eight years of data collected on a monthly basis. Each month, from August 2012 to June 2019, we considered all possible pairs among the 45 studied adult females when we had collected at least 1 focal sample (for grooming and aggression) or recorded at least one spatial association for each female of the pair. The final monthly data set contained a large majority of zeros; we therefore pooled these data together to improve our statistical models. We used Generalized Linear Models (GLM; proc GENMOD, SAS Studio) with a negative binomial distribution to study the relationships between time spent grooming (in seconds), number of aggression, and number of associations along with a set of explanatory variables. We considered as an offset the total time of observation or the total number of scans performed during the study period on each female of the pair, to adjust for variation in sampling effort. We considered as explanatory variables: the difference in social rank between the two females of the pair (class variable with three modalities: no rank difference; rank difference of one –corresponding to the difference between low- and mid-ranking females or between high- and mid-ranking females; and rank difference of two –corresponding to the difference between low- and high-ranking females); the absolute difference in age between the two females of the pair (continuous variable) and their kinship (class variable with three modalities: PHS, MHS and NK). We further considered the interaction between difference in age and kinship to control for possible combined effects. We kept full models as final models excluding only the interaction when not significant.

#### Genetic and facial similarities

To study the relationship between genetic relatedness and facial similarity (considered as a distance) calculated from pairs of photographs taken on 38 adult females with at least both parents known, we used General Linear Mixed Models (LMM; proc GLIMMIX, SAS Studio) with face distance as a response variable and the following explanatory variables. We considered the genetic relatedness of each pair (continuous variable ranging from 0.016 and 0.656) and the absolute difference in age between the two females of the pair (continuous variable). We further considered the interaction between difference in age and genetic relatedness to control for possible combined effects. We kept the full model as final model excluding only the interaction if not significant.

#### Kinship and facial similarity

To study the relationship between kinship and facial similarity (considered as a distance) calculated from pairs of photographs taken on 45 adult and on 16 juvenile females, we used General Linear Mixed Models (LMM; proc GLIMMIX, SAS Studio) with face distance as a response variable and the following explanatory variables. We considered the kinship of each pair (class variable with three modalities: PHS, MHS and NK) and the absolute difference in age between the two females of the pair (continuous variable). We further considered the interaction between difference in age and kinship to control for possible combined effects. Because we found a significant effect of this interaction on face distance in both adult and juvenile females, we repeated our analyses across different age differences to determine when this interaction is no longer significant (see results). We kept the full models as final models excluding only the interaction when not significant.

## Acknowledgments

We are grateful to past and present field assistants of the Mandrillus Project who collect daily behavioural data on the study population. This study was funded by several grants that allowed long-term collection of data: Deutsche Forschungsgemeinschaft (DFG, KA 1082-20-1) to PMK and MJEC, SEEG Lékédi (INEE-CNRS) to MJEC, and Agence Nationale de la Recherche (ANR SLEEP 17-CE02-0002) to MJEC. We are grateful to the Wildlife Reserves of Singapore and the Zoo of Granby and to the Primatological Centre at CIRMF (Gabon) for providing pictures of their mandrills. We further thank the SODEPAL-COMILOG society (ERAMET group) for their long-term logistical support and contribution to the Mandrillus Project. This study was approved by an authorization from the CENAREST institute (permit number: AR0060/18/MESRS/CENAREST/CG/CST/CSAR). This is a Project Mandrillus publication number # and ISEM 2019-##-SUD.

