## Supplementary material for "Same father, same face: deep-learning reveals paternally-derived signalling of kinship in a wild primate": SI

**SUPPLEMENTARY INFORMATION**

I. **Supplementary Figures**

**Supplementary Figure 1. Sexual dimorphism in mandrills and male’s age categories.**

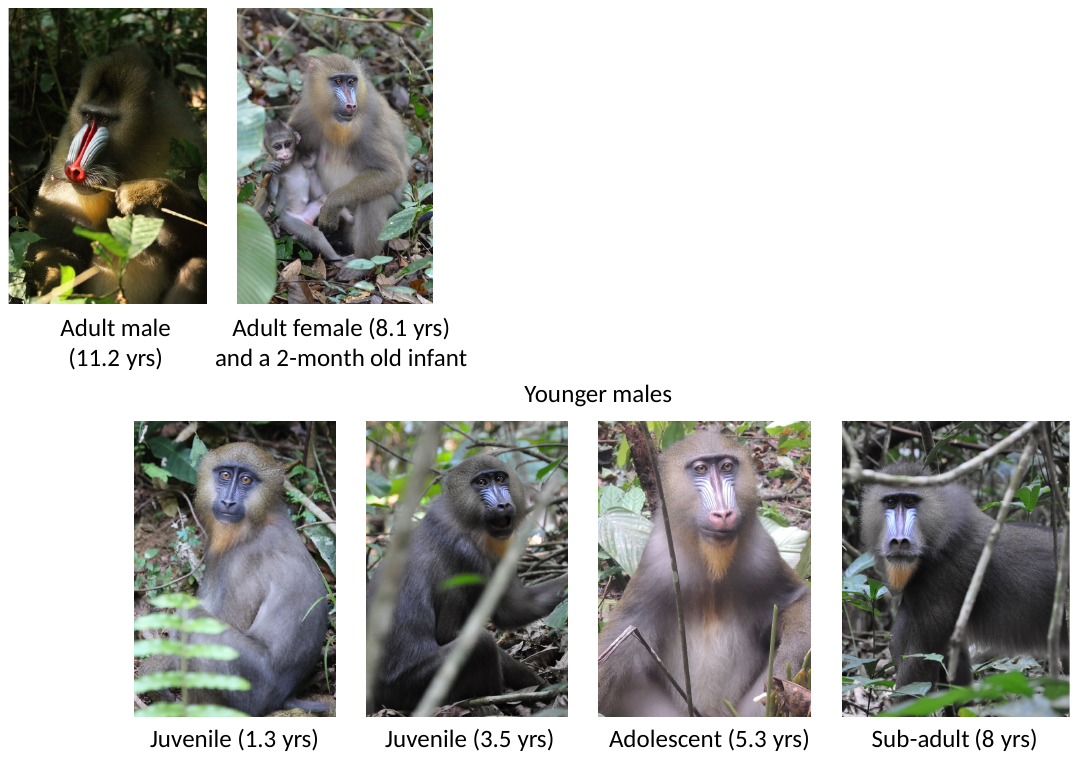

**Supplementary Figure 2. Learning curve in the face identification task.** Evolution of accuracy for the training set (f-Q1234-12-Tr; in red) and validation set (f-Q1234-12-V; in green) during a typical run. The accuracy stabilized after 15 epochs around 91% for the validation set; the run thus automatically stopped after 18 epochs (based on a stabilization of the loss). The small difference between the training accuracy and the validation accuracy indicates a limited or no effect of overfitting.

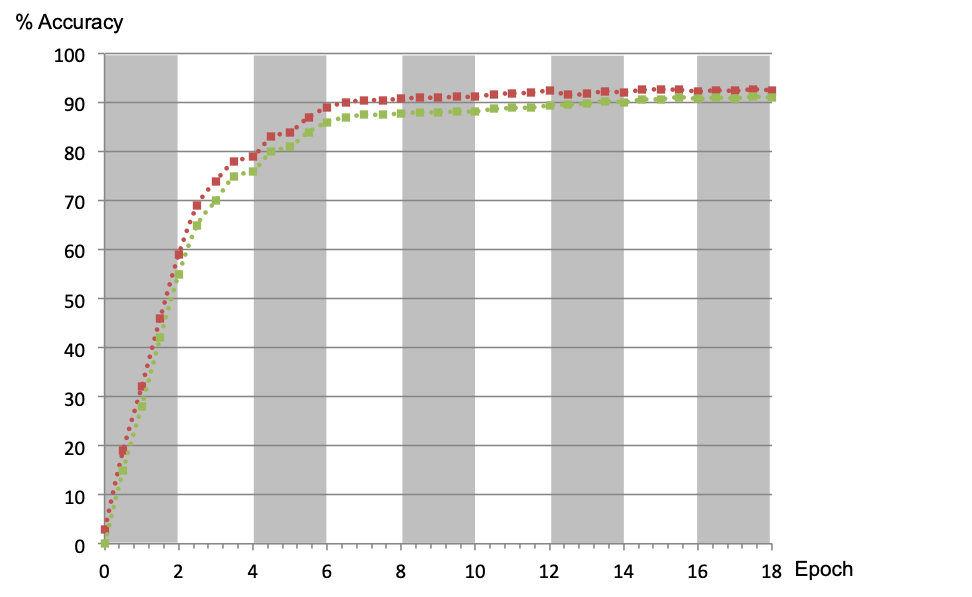

II. **Supplementary Tables**

**Supplementary Table 1. Sample sizes and facial distances (mean and SD) across pairs of females.** For comparison, we included all pairs of photographs taken, at different ages, on the same female.

|  |  | Number of pairs of females  (pairs of pictures) | Mean facial distance  (SD) |
| --- | --- | --- | --- |
| Juvenile females | PHS | 16 (111) | 12.89 (2.13) |
|  | MHS | 5 (181) | 15.80 (1.0) |
|  | NK | 27 (1297) | 15.07 (2.27) |
|  | Identical females^1^ | 53 (2087) | 10.34 (2.66) |
| Adult females | PHS | 50 (2219) | 12.83 (2.17) |
|  | MHS | 30 (1564) | 12.94 (2.08) |
|  | NK | 79 (3209) | 13.83 (1.89) |
|  | Identical females^2^ | 18 (2071) | 9.64 (3.37) |

^1^Maximum difference in age: 6.9 yrs

^2^Maximum difference in age: 2.0 yrs

**Supplementary Table 2. Characteristics of image sets**. The name of image sets is coded as followed: whether the dataset is for the adult female (f) or juvenile (j) analysis - the level of image quality (e.g., Q1234 = all qualities retained) - the minimal number of images per class before the training/validation partition (either 12 or 25) - the use of the image sets (Tr: training, V: Validation, Te: Test). Wild: wild population, ad.: adult, fem.: female, #: number of.

| **Set name** | **Sex & age categories** | **# images** | **# classes** | **Mean image/class** |
| --- | --- | --- | --- | --- |
| f-Q1234-12-Tr | all but ad. fem. from Wild | 8429 | 158 | 53.3 |
| f-Q1234-25-Tr | all but ad. fem. from Wild | 8218 | 141 | 58.3 |
| f-Q234-12-Tr | all but ad. fem. from Wild | 7914 | 132 | 60.0 |
| f-Q234-25-Tr | all but ad. fem. from Wild | 7386 | 108 | 68.4 |
| f-Q1234-12-V | all but ad. fem. from Wild | 316 | 158 | 2 |
| f-Q1234-25-V | all but ad. fem. from Wild | 182 | 141 | 2 |
| f-Q234-12-V | all but ad. fem. from Wild | 264 | 132 | 2 |
| f-Q234-25-V | all but ad. fem. from Wild | 216 | 108 | 2 |
| f-Q4-Te | ad. fem. from Wild | 421 | 55 | 7.6 |
| j-Q1234-12-Tr | all but juv. from Wild | 13772 | 202 | 68.2 |
| j-Q1234-25-Tr | all but juv. from Wild | 11901 | 155 | 76.8 |
| j-Q234-12-Tr | all but juv. from Wild | 13155 | 200 | 65.7 |
| j-Q234-25-Tr | all but juv. from Wild | 11548 | 155 | 74.5 |
| j-Q1234-12-V | all but juv. from Wild | 404 | 202 | 2 |
| j-Q1234-25-V | all but juv. from Wild | 310 | 155 | 2 |
| j-Q234-12-V | all but juv. from Wild | 400 | 200 | 2 |
| j-Q234-25-V | all but juv. from Wild | 310 | 155 | 2 |
| j-Q1234-Te | all but juv. from Wild | 472 | 50 | 9.4 |
| j-Q34-Te | juv. from Wild | 383 | 50 | 7.6 |

**Supplementary Table 3. Performance of VGG-Mandrill in face identification.** The accuracy (± sem) is averaged over 5 runs. Highest accuracy in bold.

| **Training set** | **Validation set** | **Accuracy ± sem** |
| --- | --- | --- |
| **Adult females** |  |  |
| **f-Q1234-12-Tr** | **f-Q1234-12-V** | **0.911 ± 0.118** |
| f-Q1234-25-Tr | f-Q1234-25-V | 0.898 ± 0.163 |
| f-Q234-12-Tr | f-Q234-12-V | 0.883 ± 0.297 |
| f-Q234-25-Tr | f-Q234-25-V | 0.867 ± 0.304 |
| **Juveniles** |  |  |
| **j-Q1234-12-Tr** | **j-Q1234-12-V** | **0.919 ± 0.321** |
| j-Q1234-25-Tr | j-Q1234-25-V | 0.906 ± 0.478 |
| j-Q234-12-Tr | j-Q234-12-V | 0.872 ± 0.677 |
| j-Q234-25-Tr | j-Q234-25-V | 0.851 ± 0.694 |

**Supplementary Table 4. Performance of face verification on the female adult test sets.** The accuracy (± sem) is averaged over 5 runs. Highest accuracy in bold.

| **Learning set** | **Layer** | **ReLU** | **Metric Learning** | **Accuracy ± sem** |
| --- | --- | --- | --- | --- |
| Q1234-12-Tr+V | *FC*1 | No | No | 0.639 ± 0.016 |
| Q1234-12-Tr+V | *FC*2 | No | No | 0.636 ± 0.014 |
| Q1234-12-Tr+V | *FC*12 | No | No | 0.655 ± 0.012 |
| Q1234-12-Tr+V | *FC*1 | Yes | No | 0.705 ± 0.018 |
| Q1234-12-Tr+V | *FC*2 | Yes | No | 0.652 ± 0.016 |
| Q1234-12-Tr+V | *FC*12 | Yes | No | 0.646 ± 0.010 |
| Q1234-12-Tr+V | *FC*1 | No | Yes | 0.679 ± 0.019 |
| Q1234-12-Tr+V | *FC*2 | No | Yes | 0.652 ± 0.017 |
| Q1234-12-Tr+V | *FC*12 | No | Yes | 0.699 ± 0.013 |
| **Q1234-12-Tr+V** | ***FC*1** | **Yes** | **Yes** | **0.834 ± 0.015** |
| Q1234-12-Tr+V | *FC*2 | Yes | Yes | 0.785 ± 0.016 |
| Q1234-12-Tr+V | *FC*12 | Yes | Yes | 0.796 ± 0.010 |
| Q234-25-Tr+V | *FC*1 | No | No | 0.601 ± 0.011 |
| Q234-25-Tr+V | *FC*2 | No | No | 0.611 ± 0.018 |
| Q234-25-Tr+V | *FC*12 | No | No | 0.612 ± 0.017 |
| Q234-25-Tr+V | *FC*1 | Yes | No | 0.680 ± 0.009 |
| Q234-25-Tr+V | *FC*2 | Yes | No | 0.662 ± 0.009 |
| Q234-25-Tr+V | *FC*12 | Yes | No | 0.659 ± 0.014 |
| Q234-25-Tr+V | *FC*1 | No | Yes | 0.648 ± 0.013 |
| Q234-25-Tr+V | *FC*2 | No | Yes | 0.655 ± 0.016 |
| Q234-25-Tr+V | *FC*12 | No | Yes | 0.656 ± 0.014 |
| Q234-25-Tr+V | *FC*1 | Yes | Yes | 0.820 ± 0.015 |
| Q234-25-Tr+V | *FC*2 | Yes | Yes | 0.776 ± 0.010 |
| Q234-25-Tr+V | *FC*12 | Yes | Yes | 0.785 ± 0.010 |

**Supplementary Table 5. Performance of face verification on the female juvenile testing sets.** The accuracy (± sem) is averaged over 5 runs. Highest accuracy in bold.

| **Learning set** | **Layer** | **ReLU** | **Metric Learning** | **Accuracy ± sem** |
| --- | --- | --- | --- | --- |
| Q1234-12-Tr+V | *FC*1 | No | No | 0.616 ± 0.008 |
| Q1234-12-Tr+V | *FC*2 | No | No | 0.603 ± 0.012 |
| Q1234-12-Tr+V | *FC*12 | No | No | 0.603 ± 0.011 |
| Q1234-12-Tr+V | *FC*1 | Yes | No | 0.740 ± 0.007 |
| Q1234-12-Tr+V | *FC*2 | Yes | No | 0.692 ± 0.018 |
| Q1234-12-Tr+V | *FC*12 | Yes | No | 0.699 ± 0.002 |
| Q1234-12-Tr+V | *FC*1 | No | Yes | 0.613 ± 0.009 |
| Q1234-12-Tr+V | *FC*2 | No | Yes | 0.634 ± 0.010 |
| Q1234-12-Tr+V | *FC*12 | No | Yes | 0.634 ± 0.011 |
| Q1234-12-Tr+V | *FC*1 | Yes | Yes | 0.868 ± 0.010 |
| Q1234-12-Tr+V | *FC*2 | Yes | Yes | 0.862 ± 0.009 |
| Q1234-12-Tr+V | *FC*12 | Yes | Yes | 0.856 ± 0.014 |
| Q234-25-Tr+V | *FC*1 | No | No | 0.645 ± 0.012 |
| Q234-25-Tr+V | *FC*2 | No | No | 0.627 ± 0.018 |
| Q234-25-Tr+V | *FC*12 | No | No | 0.619 ± 0.013 |
| Q234-25-Tr+V | *FC*1 | Yes | No | 0.769 ± 0.010 |
| Q234-25-Tr+V | *FC*2 | Yes | No | 0.704 ± 0.008 |
| Q234-25-Tr+V | *FC*12 | Yes | No | 0.715 ± 0.005 |
| Q234-25-Tr+V | *FC*1 | No | Yes | 0.641 ± 0.009 |
| Q234-25-Tr+V | *FC*2 | No | Yes | 0.672 ± 0.012 |
| Q234-25-Tr+V | *FC*12 | No | Yes | 0.679 ± 0.009 |
| **Q234-25-Tr+V** | ***FC*1** | **Yes** | **Yes** | **0.897 ± 0.010** |
| Q234-25-Tr+V | *FC*2 | Yes | Yes | 0.855 ± 0.013 |
| Q234-25-Tr+V | *FC*12 | Yes | Yes | 0.866 ± 0.009 |

III. **Supplementary Methods**

**Image database and pre-processing**

The mandrill face database includes ~16k images representing 276 different mandrills originating from the wild studied population (Gabon; 12.9k images), the semi-captive population of the Centre International de Recherche Medicale de Franceville (CIRMF, Gabon; 2.7k images) and other sources (internet, the Wildlife Reserves of Singapore, Zoo of Grandy; 0.4k images). Images from Gabon (wild and CIRMF populations) were taken between 2012 and 2018, using different camera models. Pictures represent individuals that are awake and passive, awake and active (i.e. feeding, grooming, vocalizing) or anesthetized during one of the bi-annual captures (represent 1.1k images). We frequently photographed active individuals using the slow burst mode of cameras, which allows to capture variation in face position and expression while avoiding identical frames. The multiple frames obtained while using the slow burst mode are hereafter referred to as “a burst-mode series”. Images were then manually oriented to align pupils horizontally, and cropped to generate square portraits centered on the nose and excluding the ears. Each portrait was manually labeled based on face position and image quality. Face position contained two levels:

- P0: the face is in profile view (approx. >30°), either from below or above, or in frontal view but significantly occluded (>50%).

- P1: the face is in frontal view (approx. <30°) and occlusion covers less than 50% of the face.

Image quality contained five levels:

- Q0: very bad quality; it impossible to recognize the individual without contextual information, even for experienced field assistants.

- Q1: poor quality; individual recognition is possible but challenging for experienced field assistants.

- Q2: medium quality; individual recognition is easy for field assistants but the portrait does not meet Q3 criteria.

- Q3: high, “passport” quality image; individual recognition is easy for field assistants, the face is in frontal view, it has a neutral expression and is not partially occluded, the image is sharp, has no shadow or lighting spot. Excludes images that meet Q4 criteria.

- Q4: a single image of a burst-mode series, which meets Q3 criteria.

We excluded P0 and Q0 images from all analyses. P1Q1234 dataset (simplified as Q1234 in the following) contained 14,8k images.

**Dataset partitions**

The Q1234 dataset represents 276 individuals belonging to one or several of the following age categories: infant, juvenile, adolescent (for males one), subadult (for males only) and adult. Because the face of an individual varies considerably between its different age categories (Supplementary Table 1), we used ind.age classes rather than individuals for the identification task, that is, we treated two ind.age classes representing the same individual as distinct and independent classes. The Q1234 dataset contains 343 ind.age classes (mean number of images per class: 50.1; range: [1,419]).

Q1234 was split into a learning set and a test set, which were different for the adult female and juvenile analyses. For the adult female analysis, the learning set included pictures of semi-captive and captive males and females of all age classes, as well as wild individuals from the studied wild population but adult females. For the juvenile analysis, the learning set was the same as above but it included wild adult females and excluded wild juveniles. Each learning set was itself split into a training set and a validation set. The validation set was used to parameterize the model for the face identification task. It contained 2 images of each class. For a correct evaluation of training performances, we ensured that none of the validation image was from a burst-mode series that also contained images present in the training set. The training set contained all other images of the learning set. Because we could reach high performances (see results) despite a large imbalance between classes in the training sets, we did not attempt to correct for this imbalance. Last, the test set included either adult females or juveniles from the wild population. To maximize the quality of resemblance measurements, we selected only Q4 images for the adult female test set. For the juvenile test set, we selected Q34 because we had insufficient Q4 images. The characteristics of the different image datasets are given in Supplementary Table 1.

**Face identification**

We trained a deep convolutional neural network (DNN) to identify individual (ind.age) mandrills as a goal to learn a deep representation of mandrill faces. We applied a transfer learning procedure (Yosinski *et al.*, 2014) by initializing the training with VGG-Face, a network that has learned to recognize 2,6k different humans from 2,6M portrait pictures. VGG-Face uses a VGG-16 architecture, which consists of 5 blocks of convolution (that filters images), nonlinear ReLU activation (that sets to zero all negative values) and max pooling (that selects a maximum locally) layers (*CB*), a flatten layer (*Fl*) that converts a 2*D* matrix feature activations into a vector, two fully connected layers (*FC*) and a softmax classification layer (*SM*). VGG-16 can thus be written as *CB*1–*CB*2–*CB*3–*CB*4–*CB*5–*Fl*–*FC*1–*FC*2–*SM*. For transfer learning, we replaced *SM* by a new layer of dimension fitting the number of classes in the new mandrill identification task, which varied with datasets. We included two dropout layers (with 50% dropout probability), one after each *FC* layer, to limit the risk of overfitting. We trained the network using a stochastic gradient descent with momentum optimizer with initial learning rate of 10^-5^ for *CB* layers and 10^-3^ for *FC* and *SM* layers. The learning rate decreased by a factor 10 every 5 epochs. Learning continued until the validation loss did not decrease further after three consecutive epochs, which required approximately 15 epochs (Supplementary Figure 2). In order to match the input size of VGG-Face, mandrill portraits were downsized to 224×224×3 prior to analyses. We set the batch size (the number of images used for optimization during one iteration) to 32. This is a small value compared to the standard practice; however, as in (Masters & Luschi, 2018) we found that a small batch size reduced overfitting significantly compared to larger sizes (64 or 128; results no shown). We limited overfitting further by using “data augmentation” (Perez & Wang, 2017). The transformations used in data augmentation were chosen to reproduce the spurious variation that could occur during the manual processing of images (i.e. cropping and alignment). Each iteration, images were shifted horizontally and vertically (by a number of pixels random selected within the range [-40 40]), rotated (range of degree: [-20 20]) and scaled (range of factor: [-0.7 1.2]). We did not flip images horizontally, assuming that some bilaterally asymmetrical features could be important for individual recognition. All deep learning analyses were ran on a single NVIDIA GeForce GTX 1080 GPU with MATLAB.

**Face verification**

After evaluating performance in face identification with the validation sets, we retrained the DNN using the full learning set to maximize the number of images. VGG-Mandrill, the newly trained DNN, was then used to extract deep feature activation vectors, a compact and informative representation of a mandrill face. The distance between feature activation vectors predicts the resemblance between images (Zhang *et al.*, 2018). In this study, we followed (Taigman *et al.*, 2014) and used a *χ*^2^ distance calculated with normalized features. Normalization was achieved by first subtracting to each activation of a vector the minimal value found for this feature across the entire learning set. This set to zero the lower boundary of the feature space (this step of the normalization is necessary *only* for testing the effect of activations before ReLU transformation; see below). Next, we divided each activation by the maximum value found for this feature across the entire learning set (or by 0.05 if the maximum value was under this threshold, to avoid division by a small number). This set to one the upper boundary of the feature space. Last, we normalized each feature vector by its L2-norm (Euclidean normalization).

Studies on face verification, the task of identifying whether two faces represent the same individual or not, performed best when learning a distance metric in the deep feature space (Ahonen *et al.*, 2006). Because the various features in the deep feature space contribute differently to predicting facial resemblance, a learned distance metric aims at finding the feature weights that optimize face verification. Following (Taigman *et al.*, 2014), we used a linear support vector machine (SVM) classifier to learn a distance metric. We randomly selected 15k pairs of images representing different individuals and 15k pairs representing same individuals, and for each pair we calculated the *χ*^2^ difference (*f*_1_[*i*]-*f*_2_[*i*])^2^/(*f*_1_[*i*]+*f*_2_[*i*]), where *f*_1_ and *f*_2_ are the normalized feature vectors of the two images in a pair and *i* the index of a feature. Then, we ran the SVM with the *χ*^2^ differences as explanatory variables and 0 (different-individual pairs) or 1 (same-individual pairs) as a response variable. The SVM output the accuracy of the face verification task as well as the weight of each feature. Weights were eventually used to calculate a weighted *χ*^2^ distance as *χ*^2^(*f*_1,_ *f*_2_) = Σ_i_*ω*_i_(*f*_1_[*i*]-*f*_2_[*i*])^2^/(*f*_1_[*i*]+*f*_2_[*i*]), where *ω* is the vector of feature weights.

**Face verification with test sets**

We evaluated the SVM trained for face verification on test tests and compared different feature spaces. Feature activations were extracted from *FC*1 (as a vector of length 4,096), *FC*2 (vector of length 4,096) or both layers (*FC*12: vector of length 8,192). We further analyzed the importance of the nonlinear transformation of the deep feature space by extracting activations either before or after the ReLU activation function. For this evaluation, we built a balanced dataset with an equal number of same-individual and different-individual pairs of portrait images, all from test sets. The number of different-individual pairs was set to match the number of same-individual pairs; different-individual pairs were randomly selected.

IV. **Supplementary Results**

**Reproductive skew and number of kin**

For each female involved in the study, we calculated an average number of maternal half-sisters (MHS) and paternal half-sisters (PHS) present in the study group at the end of the study, as follows. “MHSmin” represents the minimum number of MHS: they are accurately known thanks to either non-ambiguous behavioural observations or genetic analyses. “MHSmax” represents the probable maximum number of MHS. MHSmax was obtained by considering that each adult female gave birth to a living offspring (of either sex) every 18 months from 4yrs-old to the end of the study. We removed known deaths from MHSmax and divided this figure by two to consider only female MHS. The number of MHS for each study female was the mean between MHSmin and MHSmax. For PHS, and based on patterns of reproductive skew and on known or estimated dates of birth in the study group, we estimated that the alpha males sired on average 60% of offspring each year. We considered that non-dominant males sired, on average 15% of offspring each year. We calculated the number of PHS based on the total number of offspring born each year, on the alpha’s male tenure and male’s presence in the group, and on whether female’s father was the alpha male during her conception or not. We obtained the following figures:

|  | **Mean number (SD)** | **Variance** |
| --- | --- | --- |
| **MHS** | 4.7 (2.5) | 6.1 |
| **PHS** | 9.9 (8.8) | 77.9 |

**Face identification**

VGG-Mandrill was able to identify mandrill faces with high accuracy and generalization capacity (i.e. limited overfitting; Supplementary Figure 2). The highest accuracy was 91.1% for the adult female learning set and 91.9% for the juvenile learning set (Supplementary Table 3). The highest accuracies were reached with the sets including the largest number of images. More precisely, a wider image set (i.e. with more classes) was better than a deeper image set (i.e. with more images per class), and maximizing the number of images was more important than maximizing their quality (adding poor quality images *increased* performance).

**Face verification**

The accuracy of face verification was evaluated on test sets directly, using either Q1234-12-Tr+V (which yielded the highest accuracy in face identification) or Q234-25-TR+V (which is more stringent regarding both image quality and the number of images per class) for training the DNN to identify faces and for learning the distance metric. The highest accuracy was 83% for the adult female test set (Supplementary Table 4), and 90% for the juvenile test set (Supplementary Table 5). The higher accuracy with the juvenile set compared to that with the adult female set is expected given that same-individual pictures in this set are on average more similar than same-individual pictures in the adult female set, which contains only a single image per burst-mode series.

We found a difference in the best parameter settings between the adult female and juvenile sets only for the type of learning set. With the adult female set, the highest accuracy was reached using the fullest learning set (Q1234-12-Tr+V), as in the face identification task. With the juvenile set, the highest accuracy was reached with the more stringent learning set (Q234-25-Tr+V), as opposed to the face identification task. Note, however, that the influence of the type of learning set is less (maximum ± 3%) than that of other parameters.

The choice of the DNN layer at which feature activations are extracted influenced the accuracy of face verification by up to ± 5%. For either test set, *FC*1 activations yielded higher performance compared to *FC*2 and *FC*12 activations. A more influencing factor is the use of a new metric learned specifically for face verification. A learned distance metric raised the accuracy by as much as 15 % (with the adult female set) and 17 % (with the juvenile set). Last, the non-linearization of activations was even more influencing. Extracting activations after a ReLU transformation boosted performance by as much as 17 % (with the adult female set) and 25 % (with the juvenile set).

Based on these results, for both the adult female and juvenile analyses we estimated resemblance between pairs of mandrill faces by:

- training the DNN to perform face identification on Q1234-12-Tr+V learning set,

- extracting the *FC*1 feature activation vectors (after ReLU transformation) of Q1234-12-Tr+V images,

- training a SVM classifier with these vectors to verify face identity and compute feature weights,

- extracting the *FC*1 feature activation vectors (after ReLU transformation) of test sets,

- calculating the weighted distance between every pairs of images in each test set,

- averaging pairwise distances for every different pairs of individuals.
